# Contribution of Signaling Partner Association to Strigolactone Receptor Selectivity

**DOI:** 10.1101/2022.07.21.500684

**Authors:** Jiming Chen, Diwakar Shukla

## Abstract

The parasitic plant witchweed, or *Striga hermonthica*, results in agricultural losses of billions of dollars per year. It perceives its host via plant hormones called strigolactones, which acts as a germination stimulant for witchweed. Strigolactone signaling involves substrate binding to the strigolactone receptor followed by substrate hydrolysis and a conformational change from an inactive, or open state, to an active, or closed state. While in the active state, the receptor associates with a signaling partner, MAX2. Recently, it was shown that this MAX2 association process acts as a strong contributor to the uniquely high signaling activity observed in *Sh*HTL7, however, it is unknown why *Sh*HTL7 has enhanced MAX2 association affinity. Using an umbrella sampling molecular dynamics approach, we characterized the association processes of *At* D14, *Sh*HTL7, a mutant of *Sh*HTL7, and *Sh*HTL6 with MAX2 homolog *Os*D3. From these results, we show that *Sh*HTL7 has an enhanced standard binding free energy of *Os*D3 compared to the other receptors. Additionally, our results suggest that the overall topology of the T2-T3 helix region is likely an important modulator of MAX2 binding. Thus, differences in MAX2 association, modulated by differences in the T2-T3 helix region, are a contributor to differences in signaling activity between different strigolactone receptors.

## Introduction

Strigolactones are a class of endogenous plant hormones involved in the regulation of shoot branching, hypocotyl elongation, and root architecture in plants. ^1–3^ Signal transduction begins with binding of strigolactone to its receptor, DWARF14 (D14). This receptor also functions as a hydrolase enzyme for the hormone, and after the substrate binds, it is hydrolyzed by the receptor.^4,5^ This hydrolysis reaction covalently modifies the receptor, promoting a conformational change of the receptor to its closed, or active state which associates with MAX2 and SMXL proteins.^4,6,7^ This signaling complex is ubiquitinated and degraded, inducing a signaling response. ^4–8^ It has also been shown that MAX2 can interact with inactive-state D14 via its C-terminal helix and inhibit substrate hydrolysis,^6^ and that conformational dynamics of MAX2 plays a role in transducing the strigolactone signal. ^7^ Despite high sequence, structure, and binding pocket conservation across different strigolactone receptors, a receptor found in the parasitic plant *Striga, Sh*HTL7, has been observed to display a picomolar EC50 toward strigolactone when measured via a germination response assay.^9^ This high sensitivity has previously been attributed to the binding pocket volume of *Sh*HTL7, ^10–12^ however, this does not account for the effects of the subsequent transformations leading to the formation of the signaling complex: substrate hydrolysis, activation, and association with MAX2.

Recently, Wang *et al*. found that measured binding affinities for naturally occurring strigolactone are similar across different HTL proteins found in *Striga*. However, *Sh*HTL7 had a higher affinity for MAX2 compared to other HTL proteins when evaluated with an *in vitro* pull-down assay.^13^ This similarity in ligand affinity suggests that the binding and hydrolysis activity are similar across HTL proteins, thus the difference in MAX2 affinity of *Sh*HTL7 is likely due to either higher likelihood of being in an active conformation or a stronger interaction with MAX2 while in the active conformation. Additionally, this study found that mutating five key residues on *Sh*HTL6 to their equivalent *Sh*HTL7 residues promoted association of the mutant *Sh*HTL6 over the wild type. ^13^ In our previous work, we determined that while *Sh*HTL7 has lower propensity for *apo* activation compared to *At* D14, enhancement of activation is fifty times greater, indicating that presence of a substrate greatly enhances its likelihood of being it its active state that allows it to form a signaling complex with MAX2. ^14,15^ Since there is a ∼10000-fold difference in signaling activity between *Sh*HTL7 and other receptors,^9^ the difference in activation enhancement by covalent modification does not completely account for the difference. Here, we aim to evaluate the contribution of D14/HTL-MAX2 binding to the high signaling activity of *Sh*HTL7 by computing the standard binding free energies of association between several strigolactone receptors and the rice homolog of MAX2, *Os*D3.

Since decoupling of the effects of activation and MAX2 affinity on formation of the strigolactone receptor (SL receptor)-MAX2 complex is difficult *in vitro*, we apply molecular dynamics (MD) simulations to characterize the association pathways of active-state *At* D14, *Sh*HTL7, a mutant of *Sh*HTL7 with five residues mutated to their equivalent *Sh*HTL6 residues, and *Sh*HTL6 with *Os*D3, the *Oryza sativa* homolog of *At* MAX2. For the *Sh*HTL7 mutant, we mutated the five residues that were shown to produce *Sh*HTL7-like association properties in *Sh*HTL6. ^13^ The structure of *At* D14 in complex with *Os*D3 is shown in Fig. 1a, and the mutation sites in wild *Sh*HTL7 and their equivalent *Sh*HTL6 residues in the *Sh*HTL7 mutant are shown in Fig. 1b and c, respectively. Using an umbrella sampling procedure developed by Gumbart *et al*.,^16,17^ we computed the standard association free energies of each D14/HTL receptor with *Os*D3. Umbrella sampling provides an efficient method of calculating a free energy profile along a particular pathway by running a series of harmonically restrained simulations at short intervals along the path. ^18^ The potential of mean force (PMF) along the pathway is then recovered using unbiasing methods such as the weighted histogram analysis method (WHAM)^19^ and the multistate Bennett acceptance ratio method (MBAR).^20^ Umbrella sampling provides an advantage over traditional long MD, particularly for large systems such as protein-protein complexes, because the presence of restraints along the pathway allows for sampling of a full transition that may be inaccessible to unbiased MD simulations due to timescale constraints. This method has previously been applied to compute association of other plant hormone signaling complexes. ^21,22^ Using this procedure, we determined that the standard binding free energy of wild-type *Sh*HTL7 is greater in magnitude than *At* D14, mutant *Sh*HTL7, and *Sh*HTL6, indicating that this association process contributes to the difference in *Sh*HTL7 sensitivity in addition to activation. Additionally, we find that the high affinity of *Sh*HTL7 for MAX2 is largely driven by significant stabilization of the lid domain upon association, rather than by specific interfacial interactions that are stronger in the *Sh*HTL7-*Os*D3 complex.

**Fig. 1.**
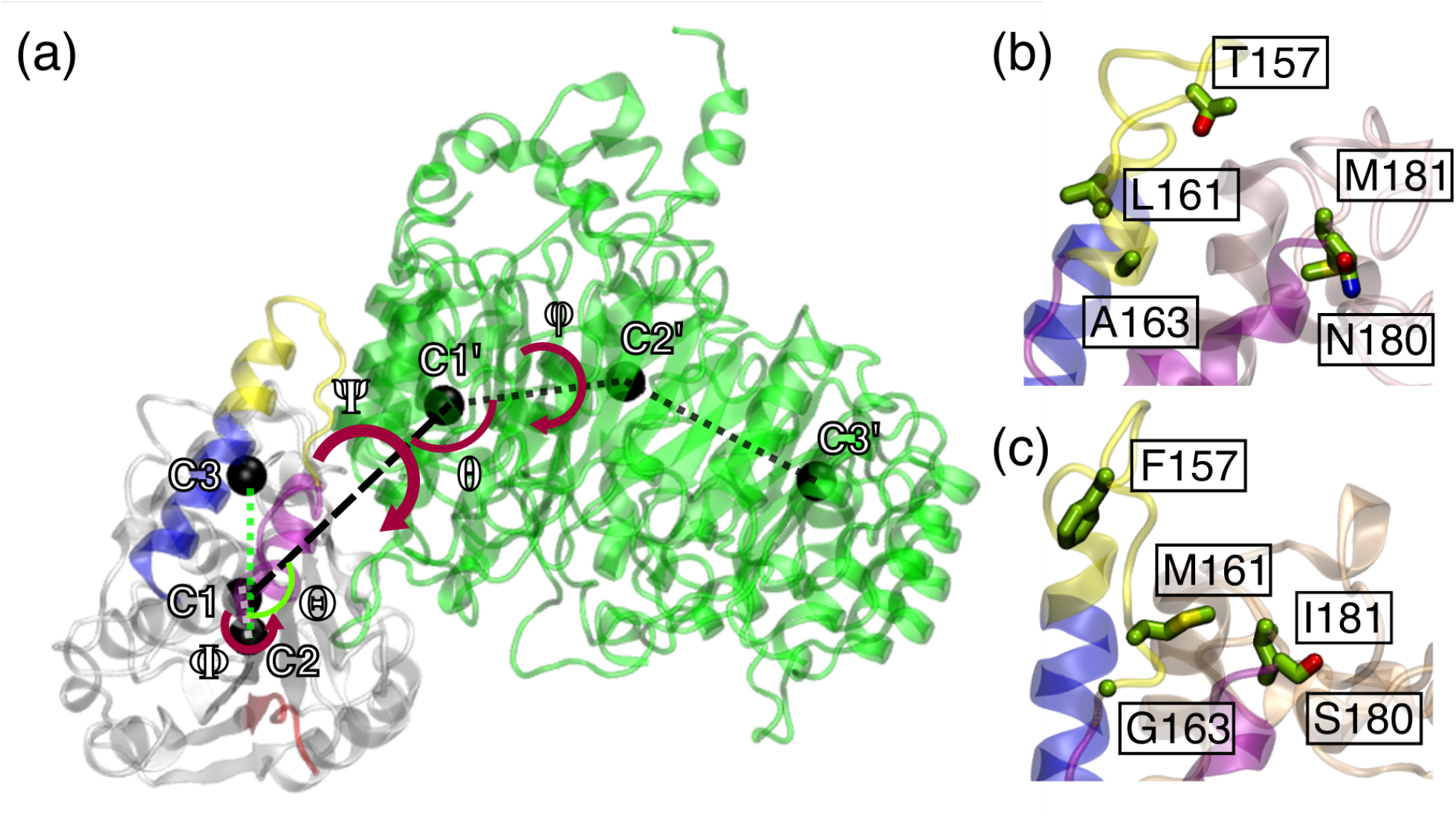
(a) Complex of *At* D14 and *Os*D3. The T1, T2, and T3 helices of *At* D14 are shown in blue, yellow, and purple, respectively, and the D-loop of *At* D14 is shown in red. *Os*MAX2 is shown in green. Centers of mass of the C1, C2, C3, C1’, C2’, and C3’ residue groups are depicted with black spheres. Restrained collective variables are defined as follows: r (separation distance): C1-C1’ distance, **Θ**: C1’-C1-C2 angle, *θ*: C1-C1’-C2’, **Φ**: C1’-C1-C2-C3 dihedral, *φ*: C1-C1’-C2’-C3’ dihedral, **Ψ**: C2-C1-C1’-C2’ dihedral. Residue groups were defined as: C1:2-263; C2:1-120, 195-263; C3:134-194; C1: 854-965; C2: 266-443; C3: 560-700 using *At* D14 numbering, with sequence-equivalent regions selected for *Sh*HTL complexes. (b)Residues mutated to *Sh*HTL6 in wild-type *Sh*HTL7 and (c) residues mutated to *Sh*HTL6 in *Sh*HTL7 mutant.

## Results

### Separation potentials of mean force similar for *At*D14, *Sh* HTL7, and *Sh* HTL6 complexes

To evaluate the association free energies of *At* D14, *Sh*HTL7, *Sh*HTL7 mutant, and *Sh*HTL6 with D3, we first applied umbrella sampling simulations to compute the potentials of mean force (PMFs) of separation. The separation PMFs for *At* D14, *Sh*HTL7 WT, and *Sh*HTL6 all had a similar depth of ∼60 kcal/mol (Fig. 2). The *Sh*HTL7 mutant had a more shallow PMF depth of ∼47 kcal/mol. These PMF depths initially appear to contradict the experimental result that *Sh*HTL7. However, it is important to note that these PMF values were computed with restraints placed on relative orientation and RMSDs of the proteins. Since the free energy associated with removing these restraints may be different for each protein complex, we computed the standard association free energies of each complex by correcting for the applied restraints using the method in Gumbart *et al*.^16,17^

**Fig. 2.**
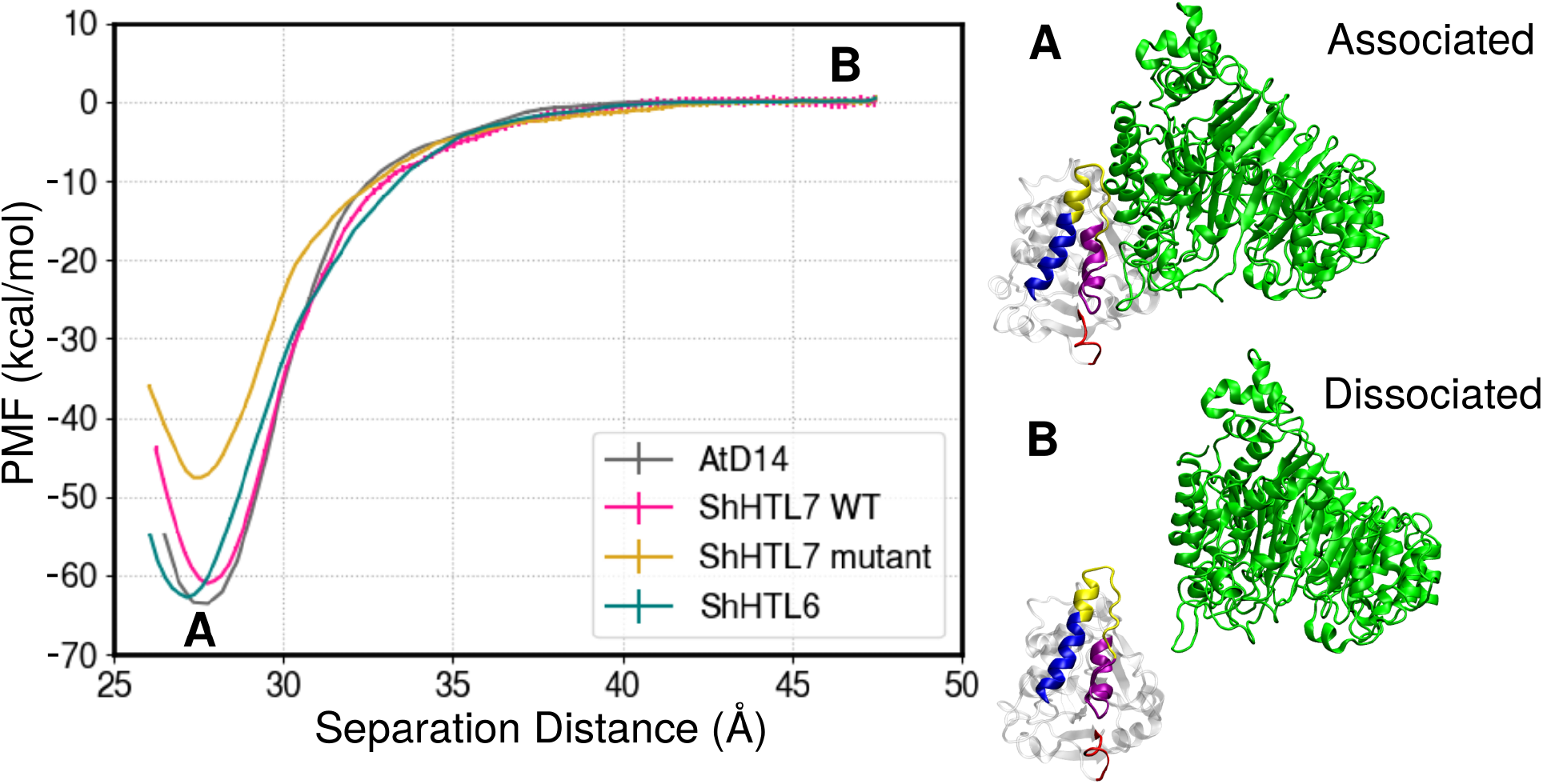
Separation potentials of mean force for *At* D14, *Sh*HTL7, an *Sh*HTL7 mutant, and *Sh*HTL6 with *Os*D3. The associated and dissociated states of *At* D14 are shown and labeled as A and B, respectively.

### Separation and RMSD terms contribute most to overall association free energy

To compute the standard free energies of binding, we performed additional umbrella sampling along all RMSD restraints and angular restraints in the associated (“site”) state. Performing additional simulations for the bulk state was unnecessary since their free energy contributions were able to be calculated analytically as detailed in the Supporting Information section on calculation of standard binding free energy. Free energy contributions of removing RMSD restraints and associated-state angular restraints are shown in Fig. 3. The exact values are shown in Table S1. For the *Sh*HTL6 complex and both *Sh*HTL7 complexes, these values are qualitatively in agreement with the findings of *Wang et al*. in that *Sh*HTL7 has a higher association affinity for MAX2 than *Sh*HTL6. ^13^ Additionally, Wang *et al*. found that mutation of five key lid helix residues on *Sh*HTL6 to equivalent *Sh*HTL7 residues enhanced association affinity, ^13^ and we show that mutation of the five residues to equivalent *Sh*HTL6 residues lowers association affinity. For all systems except the *Sh*HTL7 mutant, the separation term (*−RT* ln(*O*^*∗*^*I*^*∗*^*C°*)), where *I*^*∗*^ is the integral of the separation PMF), contributed most to the overall association free energy. Other significant (*>*10 kcal/mol) contributors to the overall association free energy were the RMSD terms. In comparison, relative orientation angle terms were small (most under ∼2 kcal/mol). When considering the contributions of RMSD restraints, the important quantity to consider is the difference between Δ*G*_RMSD_ in the bulk (unbound) and site (bound) configurations, or Δ*G*_RMSD, bulk_ *−* Δ*G*_RMSD, site_ for pair of RMSD terms: bulk and site SL receptor RMSD (labelled SL RMSD in Fig. 3) and D3 RMSD. While the differences between bulk and site Δ*G*_D3 RMSD_ show low variation across protein complexes, Δ*G*_SL RMSD_ varies widely between complexes and is an important driving force responsible for the differences in the standard binding free energies. For further investigation of the drivers of the differences in standard association free energy, we evaluated the contributions of top interacting interfacial residue pairs as well as dynamics of the SL receptors.

**Fig. 3.**
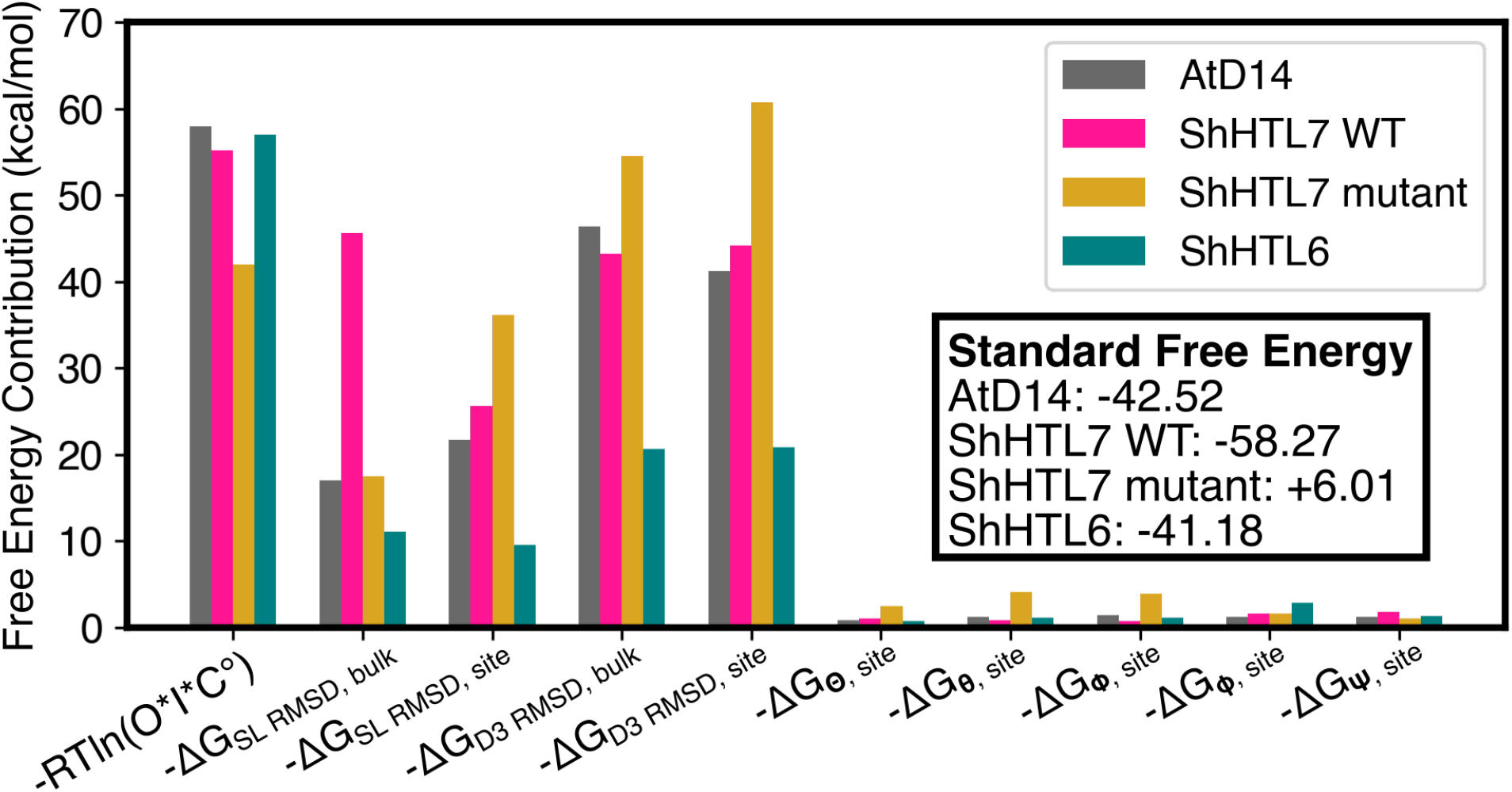
Contributions of separation (*−RT* ln(*O*^*∗*^*I*^*∗*^*C°*)), RMSD restraint terms, and angular restraint terms to overall binding free energy. For RMSD restraint terms, “bulk” refers to the D14/HTL-D3 dissociated state, and “site” refers to the D14/HTL-D3 associated state.

### Top interacting contacts show similar pair interaction energies across SL receptor-D3 complexes

To determine the key interacting residue pairs that promote receptor-D3 association, we employed pair interaction calculations in the gRINN package. ^23^ Briefly, this entailed identifying a set of frequently contacting residues on the interface between the SL receptor and *Os*D3 and computing the average potential energy of each pair interaction over the length of simulation trajectories. Pair interaction energies of the interfacial residue pairs with the top ten highest interaction energies are shown in Fig. 4a,c,e,g. Residues involved in the top five pairs with highest interaction energies are shown in Fig. 4b,d,f,h. Simulation trajectories from bound umbrella sampling windows were used for this calculation. For all four complexes, the top pair was the R702 residue on *Os*D3 forming a salt bridge with an aspartate residue on the SL receptor, D52 on *At* D14 and D29 on *Sh*HTL7 and *Sh*HTL6. For all of the *Sh*HTL-*Os*D3 complexes, the next strongest pair interaction was K34-E700. The pair interaction value for pair of residues varied from ∼50 kcal/mol to ∼80 kcal/mol, indicating that while this interaction remains a key driver of association in the different HTL receptors, the relative positioning of these interfacial residues differs slightly between complexes. Notably, the magnitudes of highest pair interaction energies of interacting residues are similar across the four protein complexes. This indicates that the differences in standard association free energy are driven by factors other than specific interfacial interactions. Since the contributions of the various restraint terms to association free energy showed that SL receptor RMSD is a strong modulator of the free energy differences, we investigated the flexibility of various regions on SL receptors in their *Os*D3 bound and unbound states.

**Fig. 4.**
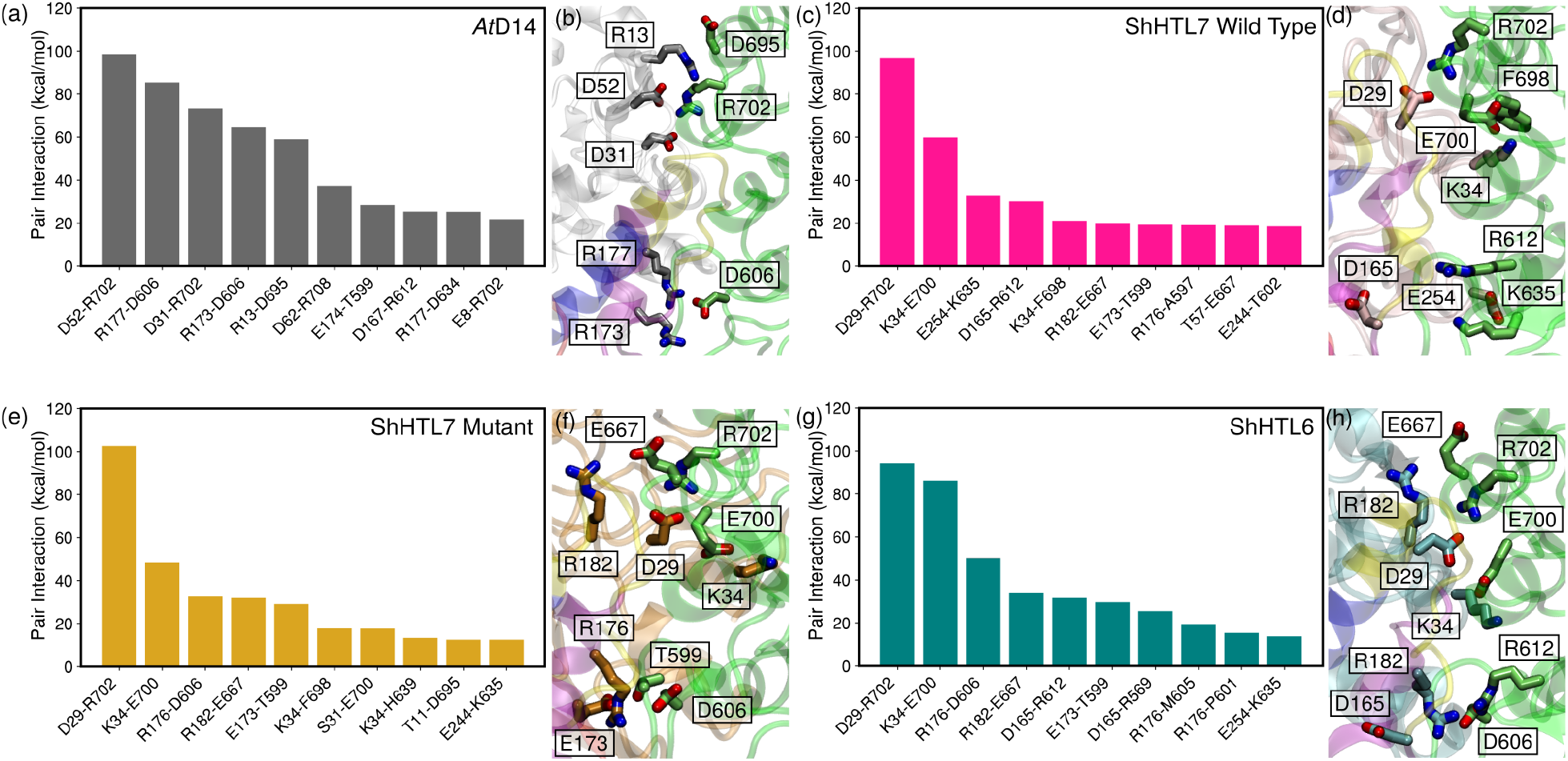
Top pair-wise residue interactions for (a,b) *At* D14, (c,d) *Sh*HTL7 Wild Type, (e,f) *Sh*HTL7 Mutant, and (g,h) *Sh*HTL6. Residue pairs with the top ten strongest attractive interactions for each protein are shown in (a,c,e,g), and residues involved in the top five interactions are shown in (b,d,f,h).

### Loss of T2-T3 region disorder upon D3 binding most prevalent in *Sh* HTL7

To evaluate why the SL receptor RMSD terms differed between the various systems, we computed crystallographic B-factors from simulations in the bound and unbound states for each complex, shown in Fig. 5. For all systems, there is a region of high B-factor, indicating greater flexibility, in the T2-T3 helix region of the lid domain. Notably, the B-factor of this region in *Sh*HTL7 is the highest of all systems in the bulk (dissociated) phase and decreases significantly in the site (associated) phase, from ∼300 Å^2^ to ∼20 Å^2^. This indicates that association with D3 stabilizes the T2 helix of wild-type *Sh*HTL7 to a greater extent than that of the other receptors considered. In *At* D14, the B-factor of the T2 helix also decreases upon D3 binding, however, the T2 helix shows a considerably lower B-factor in the unbound state (∼100 Å^2^) compared to that seen in *Sh*HTL7. This indicates that the stabilization of the T2 helix provides less contribution to the overall free energy of association, which is consistent with the lower calculated contribution of the receptor RMSD term. The *Sh*HTL7 mutant, similarly to wild type *Sh*HTL7, shows high flexibility in the T2 region, however, the B-factors of this region remain high (∼100-150 Å^2^) upon association with D3, indicating that D3 association provides less stabilization of this region in mutant *Sh*HTL7 than in wild type *Sh*HTL7. Finally, *Sh*HTL6 displays a lower B-factor in the T2 helix region than either wild-type or mutant *Sh*HTL7 in the unbound state. The T2-T3 loop in *Sh*HTL6 shows a significant decrease in B-factor upon D3 binding. This is also observed in wild-type *Sh*HTL7, thus is it unlikely that this region is a key factor for the high overall association free energy observed in *Sh*HTL7.

**Fig. 5.**
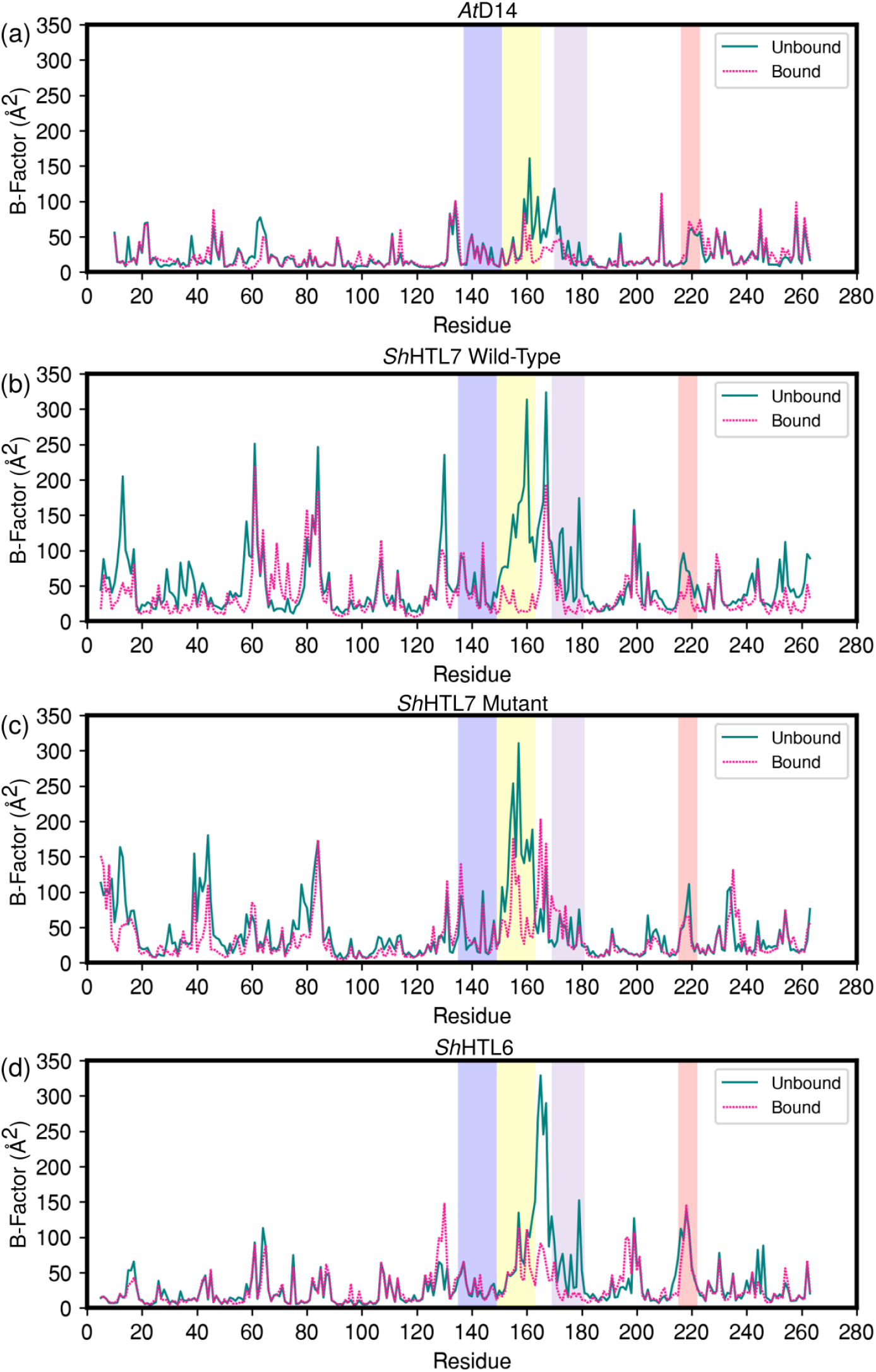
B-factors calculated in the D3-bound and unbound states for (a) *At* D14, (b) *Sh*HTL7 wild-type, (c) *Sh*HTL7 mutant, and (d) *Sh*HTL6. The T1, T2, and T3 helices are highlighted in blue, yellow, and purple, respectively. The D-loop is highlighted in red.

In all four SL receptor-D3 complexes, the unfolded portion of the T2 helix and the loop between the T2 and T3 helices form a predominantly hydrophobic interface with D3, shown in Fig. 6. Hydrophobic patches at protein-protein interfaces have previously been shown to function as important recognition sites for various protein-protein interactions. ^24–27^ A sequence alignment of *At* D14, *Sh*HTL7, and *Sh*HTL6 shows a high degree of hydrophobicity in the portion of the T2 helix that is unfolded in the active state, along with a portion of the loop between the T2 and T3 helices (Fig. S1). Specifically, the region from G158/156 on the helix to D167/165 on the T2/T3 loop is entirely made up of hydrophobic residues, with the exception of T157 on wild-type *Sh*HTL7. The conservation of this hydrophobicity suggests that the unfolded portion of the T2 helix may function as a key recognition site for association with MAX2/D3.

**Fig. 6.**
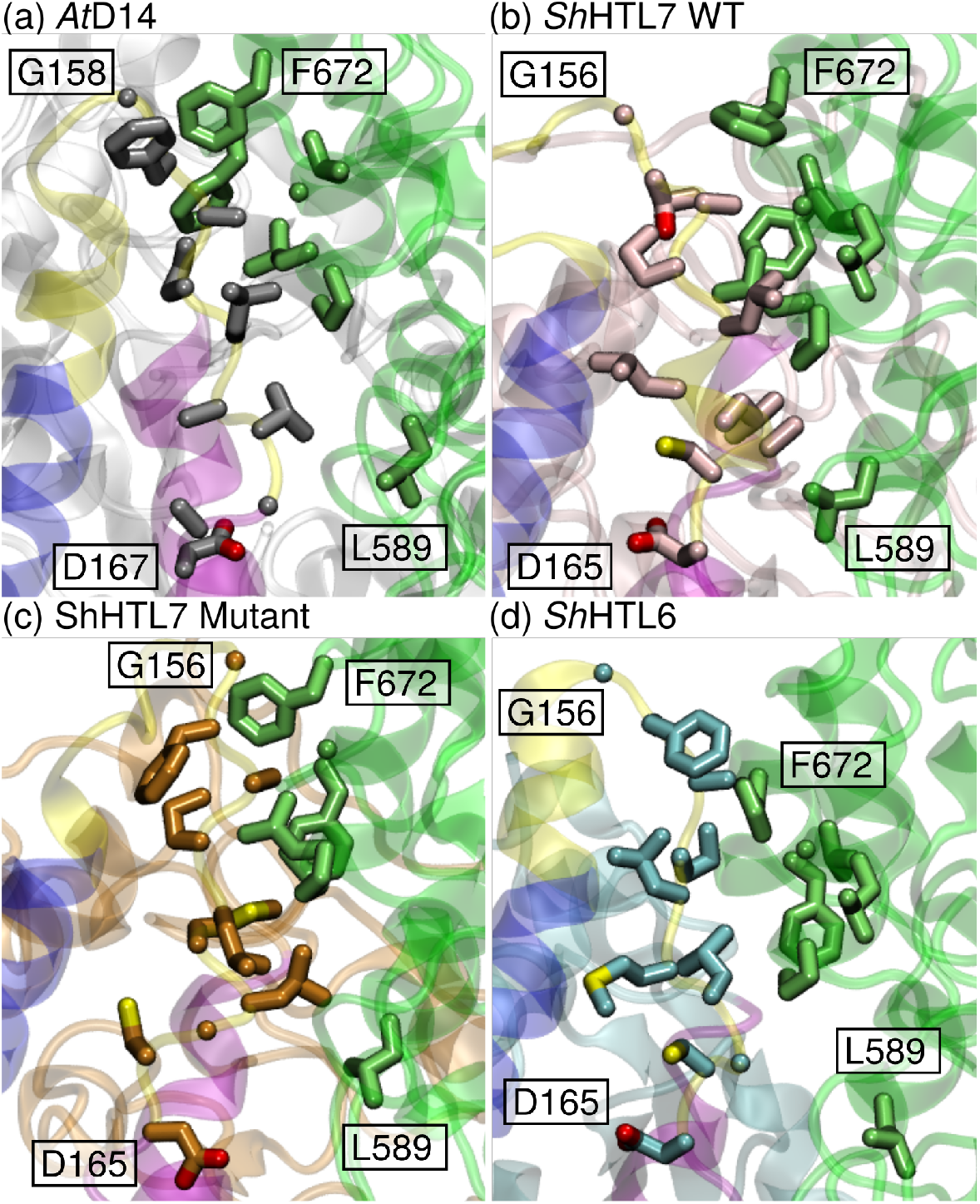
The predominantly hydrophobic interface between *Os*D3 and the unfolded T2 helix of (a) *At* D14, (b) *Sh*HTL7 wild-type, (c) *Sh*HTL7 mutant, and (d) *Sh*HTL6. The T1, T2, T3 helices of the SL receptors are shown in blue, yellow, and purple, and *Os*D3 is shown in green.

## Discussion

Using an umbrella sampling procedure, we have computed the standard association free energies of *At* D14, wild type *Sh*HTL7, mutant *Sh*HTL7, and *Sh*HTL6 with *Os*MAX2. By evaluating the contributions of various restraint terms to the overall association free energy, we have determined that differences in stabilization of the T2 helix region upon association is key to determining selectivity in *Os*D3 binding between the different receptors. Additionally, strengths of top residue pair interactions are similar across the four protein complexes, which further suggests that selectivity in association is driven by overall flexibility of the receptors, particularly in the T2-T3 helix region, rather than specific interactions with *Os*D3. Since the interfacial residues with the highest pair interaction energies are similar across the four systems evaluated and are close in pair interaction energy, our results suggest that the selectivity is modulated by conformational dynamics of the complex, specifically the T2 helix region, rather than specific interactions at the interface. This is corroborated by the magnitude of SL receptor RMSD terms during the standard binding free energy calculation. It is also corroborated by the B-factors of the four different SL receptors showing differences in T2 helix flexibility in unbound and bound states across the four receptors.

The higher association *Os*D3 association affinity of *Sh*HTL7 compared to the other receptors is consistent with the results of Wang *et al*. showing that *Sh*HTL7 shows more binding activity toward *At* MAX2 compared to other *Sh*HTL proteins.^13^ Additionally, loss of *Os*D3 association affinity when five residues in the T2-T3 helix region of *Sh*HTL7 (T157, L161, A163, N180, M181) were mutated to their equivalent *Sh*HTL6 residues (F157, M161, G163, I181, S180) suggests that this region is important for the association process. This is also consistent with the result from Wang *et al*. that mutation of these five residues in *Sh*HTL6 to the equivalent *Sh*HTL7 residues enhanced association with *At* MAX2.^13^ Notably, all of these mutations are to residues with similar chemical properties, i.e. polar to polar or hydrophobic to hydrophobic, with the exception of T157 *Sh*HTL7 and F157 in *Sh*HTL6. While this in principle could suggest that T157 in *Sh*HTL7 forms an additional hydrogen bonding interaction with MAX2 that is not present when a phenylalanine at the same position, this is not supported by our residue pair interaction strength calculations showing T157 is not involved in any of the strongest residue pair interactions at the interface. However, T157 in *Sh*HTL7 is only polar residue between G156 (*Sh*HTLs)/G158 (*At* D14) and D165 (*Sh*HTLs)/D167 (*At* D14), suggesting that it may play a role in optimizing the geometry of the region for MAX2/D3 recognition. Additionally it is also possible that this region modulates an induced fit mechanism of binding in which a conformational change is couple with association. This has been observed in other protein systems with disordered regions. ^28–30^ Several previous studies have identified this T157 residue as a important modulator of substrate binding in *Sh*HTL7. ^31–34^ Our analysis suggests that in addition to enhancing substrate binding, this residue could play a role in enhancing signaling partner association by modulating the geometry of the T2/T3 region of the receptor. The idea that the T2-T3 region in strigolactone is a key MAX2/D3 recognition region is additionally supported by the observations that although standard association free energies differ between systems, the top pair interaction energies are fairly consistent and that the RMSD of this region is a large contributor to differences in association free energy between the complexes studied.

In our previous work, we determined that *Sh*HTL7 has several advantages in substrate recognition and receptor activation that likely contribute to its high strigolactone sensitivity.^14,15,31,35^ Here, we show that signaling partner association also contributes to the high signaling ability of *Sh*HTL7. This indicates that the high MAX2 binding affinity and subsequent high sensitivity of *Sh*HTL7 is not the result of any single step in the strigolactone perception pathway on its own, but rather the combined effects of all steps. This idea also has implications in the design of small-molecule modulators to control witchweed by targeting this signaling pathway. Previous methods that have been proposed include direct inhibition of strigolactone receptors^36,37^ and induction of suicidal germination of using strigolactone mimics.^33,38,39^ Based on the finding that MAX2 association is an important contributor to the high signaling ability of *Sh*HTL7, another possible strategy is inhibition of MAX2 association. Small-molecule inhibitors of protein-protein interactions have been studied in the context of drug discovery.^40,41^ Given that differences in the T2/T3 helix region may be a key driver of selectivity in MAX2 association, small molecules that bind to this region could be a potent and selective inhibitor of witchweed germination.

## Methods

### System preparation

The associated structure of *At* D14 and *Os*D3 was obtained from PDB code 5HZG.^4^ Active state structures for other systems were constructed by homology model using Modeller. ^42–44^ Each complex was solvated in a TIP3P water box with a 0.15 M NaCl concentration. The AMBER ff14SB force field was used for proteins. Each system was minimized using a conjugate gradient descent method followed by heating to 0 to 300K and 5 ns equilibration. Temperature was maintained at 300K using the Langevin thermostat,^45^ and pressure was maintained at 1.0 bar using the Langevin barostat.^46^ Bonds to hydrogen were constrained using the SHAKE algorithm. Long-range electrostatics were computed using the particle mesh ewald algorithm.^47^ All production runs were performed using NAMD 2.14. ^48^

### Umbrella sampling protocol

A restrained umbrella sampling procedure was used to generate the separation PMFs. Harmonic restraints were placed on five orientational angles (shown in Fig. 1) and RMSDs of both proteins in each system as described in Gumbart *et al*.^16,17^ The average values of each orientational restraint over a 1 ns unbiased simulation were used to determine centers of orientational restraints. Restraint centers and force constants for each system are shown in Table S2. After the equilibration, initial structures for the umbrella sampling procedure were generated using a steered molecular dynamics simulations with orientation and RMSD restraints applied. Window centers were set at 0.33Å intervals. In total, 64 windows with centers ranging from 26Å to 46Å C1-C1’ separation distance were used for the umbrella sampling procedure. Simulation time per umbrella sampling window for each protein complex is provided in Tables S3-6.

To compute the free energy contributions of the orientation and RMSD restraints when calculating the overall association free energies, further umbrella sampling simulations were performed on each restraint. Initial structures for these simulations were obtained from adaptive biasing force (ABF) simulations.^49,50^ ABF simulations were initiated from bound umbrella sampling windows, defined as windows closest to the minimum value of the separation PMF, and were performed along both RMSD restraints and all orientational restraints. In addition, ABF simulations were also initiated from unbound umbrella sampling windows to obtain structures to calculate the PMF of removing the RMSD restraints in the unbound state. PMFS of these additional umbrella sampling simulations are shown in Fig. S2-3. Free energy contributions of angular and orientational restraints were calculated analytically, thus no additional simulations were performed for these restraints. All PMFs were calculated from umbrella sampling simulations using the multi-state Bennett acceptance ratio (MBAR) method, implemented in pyMBAR.^20^

### Free energy calculations

Overall free energies were calculated using:

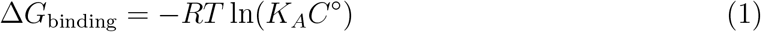

where *C*° is a standard concentration value of 1/1661 Å^3^. *K*_*A*_ was calculated using an integral of the separation PMF with corrections obtained from ensemble averages of orientation and RMSD restraints. Full details for this procedure are provided in the Supporting Information, Calculation of standard binding free energy.

### Pair interaction calculations

Pair interaction energies were computed using the pairInteraction interface in NAMD 2.14.^48^ Calculations were performed on all bound state trajectories, defined as umbrella sampling windows 5-7 for all complexes. First, a set of frequently interacting pairs were identified by calculating pairwise distances between SL receptor and D3 residues at the binding interface and selecting all pairs within a cutoff inter-residue distance of 12 Å. The average of the pair interaction energy for each pair across all simulation frames in bound windows was subsequently calculated.

### B-factor calculations

To compute B-factors, root mean square fluctuation (RMSF) were first calculated using the CPPTRAJ software.^51^ B-factors were computed using the formula:

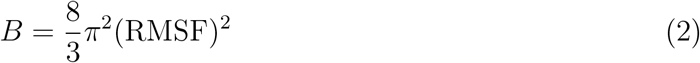

## Data and code availability

In-house code to prepare and analyze simulations is available at http://github.com/ShuklaGroup/Strigolactone-Association.

## Acknowledgments

This research was part of the Blue Waters sustained-petascale computing project, which was supported by the National Science Foundation (Award Nos. OCI-0725070 and ACI-1238993), the State of Illinois, and, as of December 2019, the National Geospatial-Intelligence Agency. Blue Waters was a joint effort of the University of Illinois at Urbana-Champaign and its National Center for Supercomputing Applications. J.C. is a member of the NIH Chemistry-Biology Interface Training Program (T32-GM136629). D.S. acknowledges support from the CAS Fellowship, Center for Advanced Studies at University of Illinois at Urbana-Champaign, and a Sloan Research Fellowship from the Alfred P. Sloan Foundation.

## Notes

### Competing Interest Statement

The authors have declared no competing interest.

