## Supplementary File for "Contribution of Signaling Partner Association to Strigolactone Receptor Selectivity"

### Supporting Information for: Contribution of Signaling Partner Association to Strigolactone Receptor Selectivity

### Contributions of Restraint Terms to Overall Free Energy

**Table 1.** Contributions of each term to overall association free energy for each of the four systems

| | *At*D14 | *Sh*HTL7 WT | *Sh*HTL7 mutant | *Sh*HTL6 |
|---|---|---|---|---|
| Separation term | -57.97 | -55.17 | -41.95 | -56.99 |
| D14/HTL RMSD, bulk | -16.97 | -45.6 | -17.45 | -11.07 |
| D14/HTL RMSD, site | -21.68 | -25.6 | -36.18 | -9.52 |
| D3 RMSD, bulk | -46.42 | -43.2 | -54.58 | -20.63 |
| D3 RMSD, site | -41.27 | -44.2 | -60.73 | -20.82 |
| $\Theta$ | -0.85 | -0.98 | -2.47 | -0.76 |
| $\theta$ | -1.21 | -0.82 | -4.11 | -1.10 |
| $\Phi$ | -1.38 | -0.72 | -3.86 | -1.11 |
| $\phi$ | -1.19 | -1.61 | -1.61 | -2.87 |
| $\Psi$ | -1.20 | -1.75 | -0.97 | -1.26 |
| Overall $\Delta G$ | -42.52 | -58.27 | 6.01 | -41.18 |

### Sequence alignment of *At*D14, *Sh*HTL7, and *Sh*HTL6

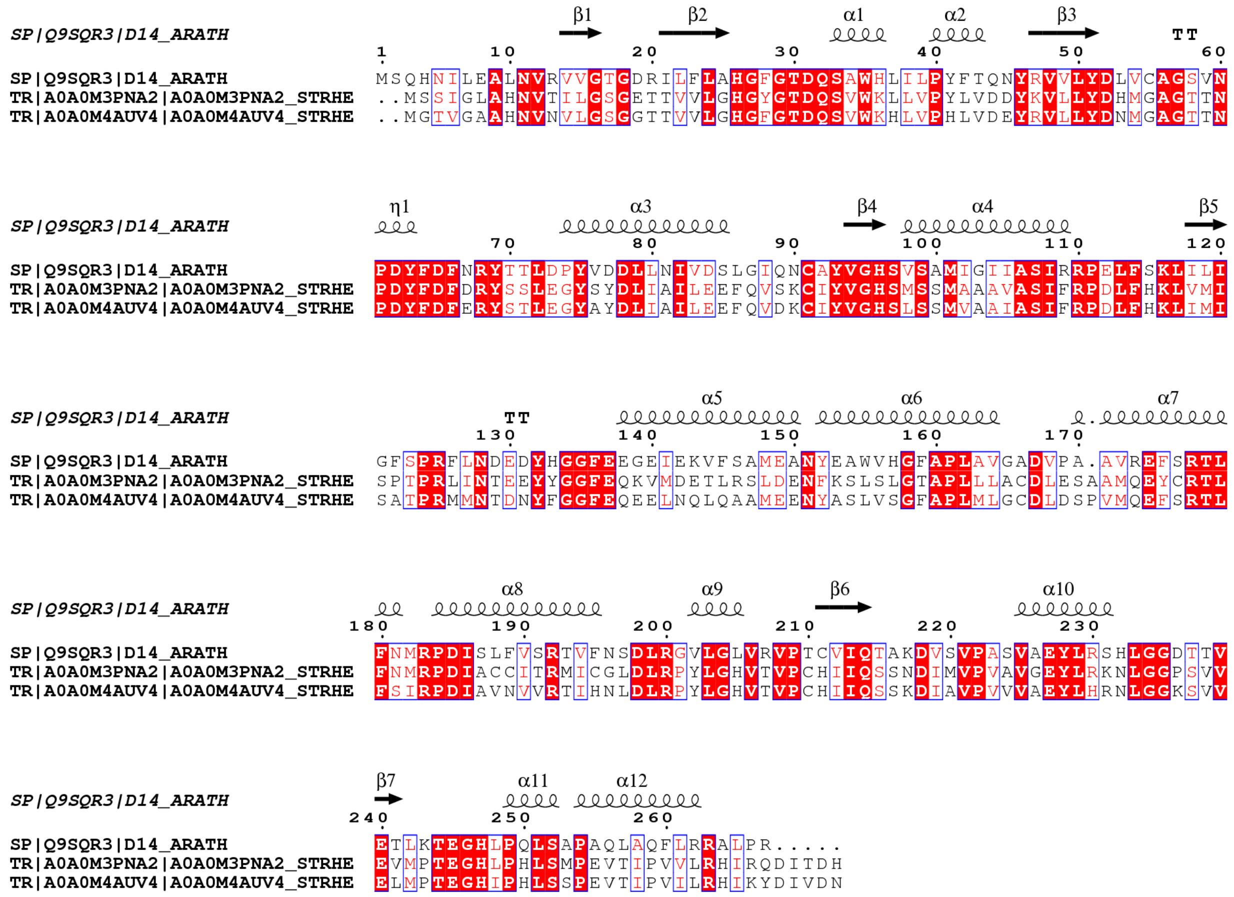


**Fig. 1.** Sequence alignment of *At*D14 (D14_ARATH), *Sh*HTL7 (A0A0M3PNA2_STRHE), and *Sh*HTL6 (A0A0M4AUV4_STRHE). Conserved residues are highlighted in red. Secondary structure elements are shown above their corresponding sequence positions.

### Restraints used during calculation of separation PMF

**Table 2.** Force constants and restraint centers for restrained collective variables during separation PMF calculation

| CV | Force constant | *At*D14 | *Sh*HTL7 WT | *Sh*HTL7 Mutant | *Sh*HTL6 |
|---|---|---|---|---|---|
| C1-C1' separation | 21.384* | - | - | - | - |
| D14/HTL7 RMSD | 20* | 0 | 0 | 0 | 0 |
| D3 RMSD | 20* | 0 | 0 | 0 | 0 |
| $\Theta$ | 10† | 84.595 | 89.191 | 83.164 | 87.724 |
| $\Phi$ | 10† | -32.386 | -38.305 | -43.600 | -37.885 |
| $\Psi$ | 10† | -14.233 | -7.533 | -30.109 | -17.096 |
| $\phi$ | 10† | -120.940 | -115.563 | -109.501 | -119.794 |
| $\theta$ | 10† | 116.422 | 105.923 | 108.683 | 116.244 |

*kcal/(mol*Å$^2$); †kcal/(mol*deg$^2$)

### Total sampling for each system

| *At*D14 | | | |
|---|---|---|---|
| Window | Replicates | Time per Replicate | Sampling for Window |
| 0 | 5 | 9.9 | 49.4 |
| 1 | 5 | 10.0 | 50.0 |
| 2 | 5 | 10.5 | 52.5 |
| 3 | 5 | 10.5 | 52.5 |
| 4 | 5 | 10.5 | 52.5 |
| 5 | 5 | 10.5 | 52.5 |
| 6 | 5 | 10.5 | 52.4 |
| 7 | 5 | 10.5 | 52.5 |
| 8 | 5 | 10.5 | 52.5 |
| 9 | 5 | 10.5 | 52.5 |
| 10 | 5 | 10.5 | 52.5 |
| 11 | 5 | 10.5 | 52.5 |
| 12 | 5 | 10.5 | 52.4 |
| 13 | 5 | 10.5 | 52.5 |
| 14 | 5 | 10.5 | 52.5 |
| 15 | 5 | 10.5 | 52.4 |
| 16 | 5 | 10.5 | 52.4 |
| 17 | 5 | 10.5 | 52.4 |
| 18 | 5 | 10.5 | 52.4 |
| 19 | 5 | 10.5 | 52.5 |
| 20 | 5 | 10.5 | 52.5 |
| 21 | 5 | 10.5 | 52.5 |
| 22 | 5 | 10.5 | 52.5 |
| 23 | 5 | 10.5 | 52.5 |
| 24 | 5 | 10.5 | 52.5 |
| 25 | 5 | 10.5 | 52.5 |
| 26 | 5 | 10.5 | 52.5 |
| 27 | 5 | 10.5 | 52.5 |
| 28 | 5 | 10.5 | 52.5 |
| 29 | 5 | 10.5 | 52.5 |
| 30 | 5 | 10.5 | 52.5 |
| 31 | 5 | 10.5 | 52.5 |
| 32 | 5 | 10.5 | 52.5 |
| 33 | 5 | 10.5 | 52.5 |
| 34 | 5 | 10.5 | 52.5 |
| 35 | 5 | 10.5 | 52.5 |
| 36 | 5 | 10.5 | 52.5 |
| 37 | 5 | 10.5 | 52.5 |
| 38 | 5 | 10.5 | 52.5 |
| 39 | 5 | 10.5 | 52.5 |

| | | | |
|---|---|---|---|
| 40 | 5 | 10.5 | 52.5 |
| 41 | 5 | 10.5 | 52.5 |
| 42 | 5 | 10.5 | 52.5 |
| 43 | 5 | 10.5 | 52.5 |
| 44 | 5 | 10.5 | 52.5 |
| 45 | 5 | 10.5 | 52.5 |
| 46 | 5 | 10.5 | 52.5 |
| 47 | 5 | 10.5 | 52.5 |
| 48 | 5 | 10.5 | 52.5 |
| 49 | 5 | 10.5 | 52.4 |
| 50 | 5 | 10.5 | 52.5 |
| 51 | 5 | 10.5 | 52.5 |
| 52 | 5 | 10.5 | 52.5 |
| 53 | 5 | 10.5 | 52.5 |
| 54 | 5 | 10.5 | 52.5 |
| 55 | 5 | 10.5 | 52.5 |
| 56 | 5 | 10.5 | 52.5 |
| 57 | 5 | 10.5 | 52.5 |
| 58 | 5 | 10.5 | 52.4 |
| 59 | 5 | 10.5 | 52.5 |
| 60 | 5 | 10.5 | 52.4 |
| 61 | 5 | 10.5 | 52.5 |
| 62 | 5 | 10.5 | 52.4 |
| 63 | 5 | 10.5 | 52.4 |

**Table 3.** Sampling Table for *At*D14

| *Sh*HTL7 Wild-Type | | | |
|---|---|---|---|
| Window | Replicates | Time per Replicate | Sampling for Window |
| 0 | 5 | 8.7 | 43.4 |
| 1 | 5 | 8.7 | 43.4 |
| 2 | 5 | 8.7 | 43.4 |
| 3 | 5 | 8.7 | 43.5 |
| 4 | 5 | 8.7 | 43.5 |
| 5 | 5 | 8.7 | 43.5 |
| 6 | 5 | 8.7 | 43.5 |
| 7 | 5 | 8.7 | 43.5 |
| 8 | 5 | 8.7 | 43.5 |
| 9 | 5 | 8.7 | 43.5 |
| 10 | 5 | 8.7 | 43.5 |
| 11 | 5 | 8.7 | 43.5 |
| 12 | 5 | 8.7 | 43.5 |
| 13 | 5 | 8.7 | 43.5 |
| 14 | 5 | 8.7 | 43.5 |
| 15 | 5 | 8.7 | 43.5 |
| 16 | 5 | 8.6 | 42.9 |
| 17 | 5 | 8.7 | 43.5 |
| 18 | 5 | 8.7 | 43.5 |
| 19 | 5 | 8.7 | 43.5 |
| 20 | 5 | 8.7 | 43.5 |
| 21 | 5 | 8.7 | 43.4 |
| 22 | 5 | 8.7 | 43.5 |
| 23 | 5 | 8.7 | 43.5 |
| 24 | 5 | 8.7 | 43.5 |
| 25 | 5 | 8.7 | 43.5 |
| 26 | 5 | 8.7 | 43.5 |
| 27 | 5 | 8.6 | 42.9 |
| 28 | 5 | 8.2 | 40.9 |
| 29 | 5 | 8.7 | 43.5 |
| 30 | 5 | 8.7 | 43.5 |
| 31 | 5 | 8.7 | 43.5 |
| 32 | 5 | 8.7 | 43.5 |
| 33 | 5 | 8.7 | 43.5 |
| 34 | 5 | 8.7 | 43.5 |
| 35 | 5 | 8.7 | 43.5 |
| 36 | 5 | 8.7 | 43.5 |
| 37 | 5 | 8.7 | 43.5 |
| 38 | 5 | 8.7 | 43.5 |
| 39 | 5 | 8.7 | 43.5 |
| 40 | 5 | 8.7 | 43.5 |
| 41 | 5 | 8.7 | 43.5 |

| | | | |
|---|---|---|---|
| 42 | 5 | 8.7 | 43.5 |
| 43 | 5 | 8.7 | 43.5 |
| 44 | 5 | 8.7 | 43.5 |
| 45 | 5 | 8.7 | 43.5 |
| 46 | 5 | 8.7 | 43.5 |
| 47 | 5 | 8.7 | 43.5 |
| 48 | 5 | 8.7 | 43.5 |
| 49 | 5 | 8.7 | 43.5 |
| 50 | 5 | 8.7 | 43.5 |
| 51 | 5 | 8.7 | 43.5 |
| 52 | 5 | 8.7 | 43.5 |
| 53 | 5 | 8.7 | 43.5 |
| 54 | 5 | 8.7 | 43.5 |
| 55 | 5 | 8.7 | 43.5 |
| 56 | 5 | 8.7 | 43.5 |
| 57 | 5 | 8.7 | 43.5 |
| 58 | 5 | 8.7 | 43.5 |
| 59 | 5 | 8.7 | 43.5 |
| 60 | 5 | 8.7 | 43.5 |
| 61 | 5 | 8.7 | 43.5 |
| 62 | 5 | 8.7 | 43.5 |
| 63 | 5 | 8.7 | 43.5 |

**Table 4.** Sampling Table for *Sh*HTL7 WT

| *Sh*HTL7 Mutant | | | |
|---|---|---|---|
| Window | Replicates | Time per Replicate | Sampling for Window |
| 0 | 5 | 5.1 | 25.4 |
| 1 | 5 | 5.1 | 25.4 |
| 2 | 5 | 5.1 | 25.4 |
| 3 | 5 | 5.1 | 25.4 |
| 4 | 5 | 5.1 | 25.4 |
| 5 | 5 | 5.1 | 25.4 |
| 6 | 5 | 5.1 | 25.4 |
| 7 | 5 | 5.1 | 25.4 |
| 8 | 5 | 5.1 | 25.4 |
| 9 | 5 | 5.1 | 25.4 |
| 10 | 5 | 5.1 | 25.4 |
| 11 | 5 | 5.1 | 25.4 |
| 12 | 5 | 5.1 | 25.4 |
| 13 | 5 | 5.1 | 25.4 |
| 14 | 5 | 5.1 | 25.4 |
| 15 | 5 | 5.1 | 25.4 |
| 16 | 5 | 5.1 | 25.4 |
| 17 | 5 | 5.1 | 25.4 |
| 18 | 5 | 5.1 | 25.5 |
| 19 | 5 | 5.1 | 25.5 |
| 20 | 5 | 5.1 | 25.4 |
| 21 | 5 | 5.1 | 25.4 |
| 22 | 5 | 5.1 | 25.5 |
| 23 | 5 | 5.1 | 25.4 |
| 24 | 5 | 5.1 | 25.5 |
| 25 | 5 | 5.1 | 25.4 |
| 26 | 5 | 5.1 | 25.4 |
| 27 | 5 | 5.1 | 25.4 |
| 28 | 5 | 5.1 | 25.4 |
| 29 | 5 | 5.1 | 25.4 |
| 30 | 5 | 5.1 | 25.4 |
| 31 | 5 | 5.1 | 25.4 |
| 32 | 5 | 5.1 | 25.4 |
| 33 | 5 | 5.1 | 25.4 |
| 34 | 5 | 5.1 | 25.4 |
| 35 | 5 | 5.1 | 25.4 |
| 36 | 5 | 5.1 | 25.4 |
| 37 | 5 | 5.1 | 25.4 |
| 38 | 5 | 5.1 | 25.4 |
| 39 | 5 | 5.1 | 25.4 |
| 40 | 5 | 5.1 | 25.4 |
| 41 | 5 | 5.1 | 25.4 |

| | | | |
|---|---|---|---|
| 42 | 5 | 5.1 | 25.4 |
| 43 | 5 | 5.1 | 25.4 |
| 44 | 5 | 5.1 | 25.5 |
| 45 | 5 | 5.1 | 25.4 |
| 46 | 5 | 5.1 | 25.4 |
| 47 | 5 | 5.1 | 25.4 |
| 48 | 5 | 5.1 | 25.4 |
| 49 | 5 | 5.1 | 25.4 |
| 50 | 5 | 5.1 | 25.4 |
| 51 | 5 | 5.1 | 25.4 |
| 52 | 5 | 5.1 | 25.4 |
| 53 | 5 | 5.1 | 25.4 |
| 54 | 5 | 5.1 | 25.4 |
| 55 | 5 | 5.1 | 25.4 |
| 56 | 5 | 5.1 | 25.4 |
| 57 | 5 | 5.1 | 25.4 |
| 58 | 5 | 5.1 | 25.4 |
| 59 | 5 | 5.1 | 25.4 |
| 60 | 5 | 5.1 | 25.4 |
| 61 | 5 | 5.1 | 25.4 |
| 62 | 5 | 5.1 | 25.4 |
| 63 | 5 | 5.1 | 25.4 |

**Table 5.** Sampling Table for *Sh*HTL7 Mutant

| *Sh*HTL6 | | | |
|---|---|---|---|
| Window | Replicates | Time per Replicate | Sampling for Window |
| 0 | 5 | 8.4 | 42.1 |
| 1 | 5 | 8.4 | 42.2 |
| 2 | 5 | 8.4 | 42.0 |
| 3 | 5 | 8.4 | 42.1 |
| 4 | 5 | 8.4 | 42.0 |
| 5 | 5 | 8.4 | 42.0 |
| 6 | 5 | 8.8 | 44.2 |
| 7 | 5 | 8.8 | 44.2 |
| 8 | 5 | 8.4 | 42.0 |
| 9 | 5 | 8.4 | 42.1 |
| 10 | 5 | 8.4 | 42.1 |
| 11 | 5 | 8.1 | 40.4 |
| 12 | 5 | 8.8 | 44.2 |
| 13 | 5 | 8.8 | 44.2 |
| 14 | 5 | 8.4 | 42.1 |
| 15 | 5 | 8.4 | 42.1 |
| 16 | 5 | 8.8 | 44.2 |
| 17 | 5 | 8.8 | 44.2 |
| 18 | 5 | 8.8 | 44.1 |
| 19 | 5 | 8.8 | 44.2 |
| 20 | 5 | 8.8 | 44.2 |
| 21 | 5 | 8.8 | 44.2 |
| 22 | 5 | 8.8 | 44.1 |
| 23 | 5 | 8.8 | 44.2 |
| 24 | 5 | 8.8 | 44.2 |
| 25 | 5 | 8.8 | 44.2 |
| 26 | 5 | 8.8 | 44.2 |
| 27 | 5 | 8.8 | 44.2 |
| 28 | 5 | 8.8 | 44.2 |
| 29 | 5 | 8.8 | 44.2 |
| 30 | 5 | 8.8 | 44.2 |
| 31 | 5 | 8.8 | 44.2 |
| 32 | 5 | 8.8 | 44.2 |
| 33 | 5 | 8.8 | 44.2 |
| 34 | 5 | 8.8 | 44.2 |
| 35 | 5 | 8.8 | 44.2 |
| 36 | 5 | 8.8 | 44.1 |
| 37 | 5 | 8.8 | 44.2 |
| 38 | 5 | 8.8 | 44.2 |
| 39 | 5 | 8.8 | 44.2 |
| 40 | 5 | 8.8 | 44.2 |
| 41 | 5 | 8.8 | 44.2 |

| | | | |
|---|---|---|---|
| 42 | 5 | 8.8 | 44.2 |
| 43 | 5 | 8.8 | 44.1 |
| 44 | 5 | 8.8 | 44.1 |
| 45 | 5 | 8.8 | 44.2 |
| 46 | 5 | 8.8 | 44.1 |
| 47 | 5 | 8.8 | 44.1 |
| 48 | 5 | 8.8 | 44.2 |
| 49 | 5 | 8.8 | 44.2 |
| 50 | 5 | 8.8 | 44.2 |
| 51 | 5 | 8.8 | 44.2 |
| 52 | 5 | 8.8 | 44.2 |
| 53 | 5 | 8.8 | 44.2 |
| 54 | 5 | 8.8 | 44.2 |
| 55 | 5 | 8.8 | 44.1 |
| 56 | 5 | 8.8 | 44.2 |
| 57 | 5 | 8.8 | 44.2 |
| 58 | 5 | 8.8 | 44.2 |
| 59 | 5 | 8.8 | 44.1 |
| 60 | 5 | 8.8 | 44.2 |
| 61 | 5 | 8.8 | 44.2 |
| 62 | 5 | 8.8 | 44.2 |
| 63 | 5 | 8.8 | 44.2 |

**Table 6.** Sampling Table for *Sh*HTL6

### PMFs of additional umbrella sampling simulations to correct for applied restraints

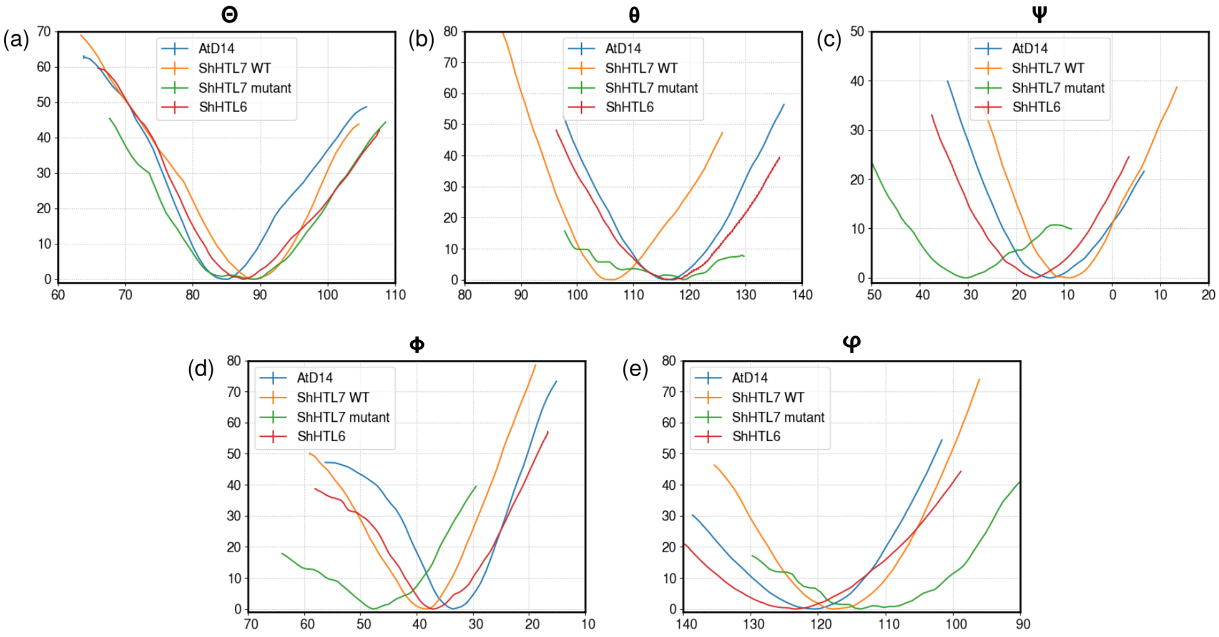


**Fig. 2.** PMFs calculated from umbrella sampling simulations of angular and orientational restraints in the bound ("site") phase of *At*D14-*Os*D3 association.

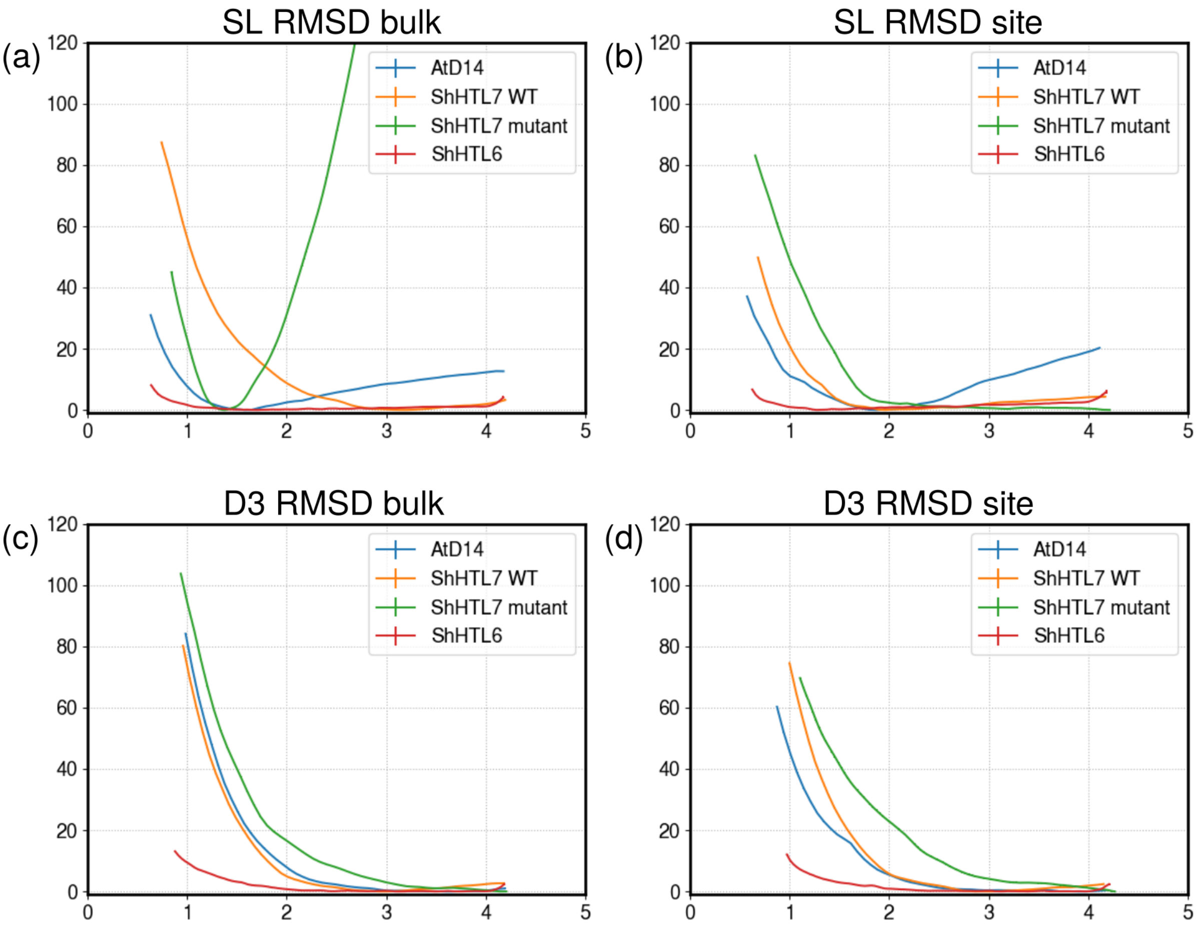


**Fig. 3.** PMFs calculated from umbrella sampling simulations of RMSD restraints in both the bound ("site") phase and unbound ("bulk") phase of *At*D14-*Os*D3 association.

### Calculation of standard binding free energy

The standard free energy of binding is calculated using the association constant:

$$\Delta G_{\text{binding}} = -RT \ln(K_A C^{\circ}) \tag{1}$$

Calculation of the association constant requires the integral of the separation PMF (Eq. 2). $W(r)$ refers to the separation PMF, and $r^*$ is the distance at which the interaction strength between the two proteins becomes equal to that of infinite separation:

$$I^* = \int_{\text{bound}} e^{-\beta[W(r)-W(r^*)]} dr \tag{2}$$

The orientational restraint contribution can be calculated using Eq. 3, where $u_a$ is the functional form of the harmonic restraints placed on the orientational variables $\theta$ and $\phi$:

$$O^* = (r^*)^2 \int_0^{\pi} \int_0^{2\pi} \sin(\theta) e^{-\beta u_a(\theta,\phi)} d\theta d\phi \tag{3}$$

The remaining portions of the $K_A$ calculation are the contributions of angular and RMSD restraints (Eq. 4):

$$K_A = O^* I^* e^{-\beta[(\Delta G^{\text{B}}_{\text{D14/HTL}} - \Delta G^{\text{S}}_{\text{D14/HTL}}) + (\Delta G^{\text{B}}_{\text{D3}} - \Delta G^{\text{S}}_{\text{D3}}) + (\Delta G^{\text{B}}_{0} - \Delta G^{\text{S}}_{0})]} \tag{4}$$

The bulk (dissociated state) angular restraint contribution can be calculated using Eq. 5, where $u_0$ is the functional form of the harmonic restraints placed on the angular variables $\Theta$, $\Phi$, and $\Psi$:

$$\Delta G^{\text{B}}_{0} = \Delta G^{\text{B}}_{\Theta} + \Delta G^{\text{B}}_{\Phi} + \Delta G^{\text{B}}_{\Psi} = -RT \ln\left[\frac{1}{8\pi^2} \int_0^{\pi} \int_0^{2\pi} \int_0^{2\pi} \sin(\Theta) e^{-\beta u_0(\Theta,\Phi,\Psi)} d\Theta d\Phi d\Psi\right] \tag{5}$$

Finally, the remaining site (associated state) angular/orientational contributions and RMSD contributions are calculated from additional simulations. The total site angular/orientational contribution is calculated using the sum of individual contributions of angular and orientational terms (Eq. 6).

$$\Delta G_0^{\mathrm{S}} = \Delta G_\Theta^{\mathrm{S}} + \Delta G_\Phi^{\mathrm{S}} + \Delta G_\Psi^{\mathrm{S}} + \Delta G_\theta^{\mathrm{S}} + \Delta G_\phi^{\mathrm{S}} \tag{6}$$

Each SL receptor RMSD term, D3 RMSD term, and angular restraint term is calculated using ensemble averages. These ensemble averages require additional umbrella sampling simulations along each restraint:

$$\Delta G_{\mathrm{D14/HTL}}^{\mathrm{S}} = RT \ln \left[ \frac{\int_S e^{-\beta[U(x)+pV(x)+u_{\mathrm{D14/HTL}}(x)]} dx}{\int_S e^{-\beta[U(x)+pV(x)]} dx} \right] \tag{7}$$

$$\Delta G_{\mathrm{D3}}^{\mathrm{S}} = RT \ln \left[ \frac{\int_S e^{-\beta[U(x)+pV(x)+u_{\mathrm{D14/HTL}}(x)+u_{\mathrm{D3}}(x)]} dx}{\int_S e^{-\beta[U(x)+pV(x)+u_{\mathrm{D14/HTL}}(x)]} dx} \right] \tag{8}$$

$$\Delta G_\Theta^{\mathrm{S}} = RT \ln \left[ \frac{\int_S e^{-\beta[U(x)+pV(x)+u_{\mathrm{D14/HTL}}(x)+u_{\mathrm{D3}}(x)+u_\Theta(x)]} dx}{\int_S e^{-\beta[U(x)+pV(x)+u_{\mathrm{D14/HTL}}(x)+u_{\mathrm{D3}}(x)]} dx} \right] \tag{9}$$

$$\Delta G_\Phi^{\mathrm{S}} = RT \ln \left[ \frac{\int_S e^{-\beta[U(x)+pV(x)+u_{\mathrm{D14/HTL}}(x)+u_{\mathrm{D3}}(x)+u_\Theta(x)+u_\Phi(x)]} dx}{\int_S e^{-\beta[U(x)+pV(x)+u_{\mathrm{D14/HTL}}(x)+u_{\mathrm{D3}}(x)+u_\Theta(x)]} dx} \right] \tag{10}$$

$$\Delta G_\Psi^{\mathrm{S}} = RT \ln \left[ \frac{\int_S e^{-\beta[U(x)+pV(x)+u_{\mathrm{D14/HTL}}(x)+u_{\mathrm{D3}}(x)+u_\Theta(x)+u_\Phi(x)+u_\Psi(x)]} dx}{\int_S e^{-\beta[U(x)+pV(x)+u_{\mathrm{D14/HTL}}(x)+u_{\mathrm{D3}}(x)+u_\Theta(x)+u_\Phi(x)]} dx} \right] \tag{11}$$

$$\Delta G_\phi^{\mathrm{S}} = RT \ln \left[ \frac{\int_S e^{-\beta[U(x)+pV(x)+u_{\mathrm{D14/HTL}}(x)+u_{\mathrm{D3}}(x)+u_\Theta(x)+u_\Phi(x)+u_\Psi(x)+u_\phi(x)]} dx}{\int_S e^{-\beta[U(x)+pV(x)+u_{\mathrm{D14/HTL}}(x)+u_{\mathrm{D3}}(x)+u_\Theta(x)+u_\Phi(x)+u_\Psi(x)]} dx} \right] \tag{12}$$

$$\Delta G_\theta^{\mathrm{S}} = RT \ln \left[ \frac{\int_S e^{-\beta[U(x)+pV(x)+u_{\mathrm{D14/HTL}}(x)+u_{\mathrm{D3}}(x)+u_\Theta(x)+u_\Phi(x)+u_\Psi(x)+u_\phi(x)+u_\theta(x)]} dx}{\int_S e^{-\beta[U(x)+pV(x)+u_{\mathrm{D14/HTL}}(x)+u_{\mathrm{D3}}(x)+u_\Theta(x)+u_\Phi(x)+u_\Psi(x)+u_\phi(x)]} dx} \right] \tag{13}$$

Using PMFs obtained from umbrella sampling simulations, each ensemble average can be calculated using the integral of the PMF along each restraint variable, for example:

$$\Delta G_{\theta}^{\mathrm{S}} \approx RT \ln \left[ \int_{S} w^{\theta}_{\mathrm{D14/HTL,\ D3},\Theta,\Phi,\Psi,\phi} \right] \tag{14}$$

In this example, $w^{\theta}_{\mathrm{D14/HTL,\ D3},\Theta,\Phi,\Psi,\phi}$ is the PMF obtained from an umbrella sampling simulation over $\theta$ with variables D14/HTL, $\mathrm{D3}, \Theta, \Phi, \Psi, \phi$ restrained.
